# Flumequine, a fluoroquinolone in disguise

**DOI:** 10.1101/2024.04.19.590246

**Authors:** Aram F. Swinkels, Egil A.J. Fischer, Lisa korving, Rafaella Christodoulou, Jaap A. Wagenaar, Aldert L. Zomer

## Abstract

Fluoroquinolone resistance in *E. coli* isolates from livestock in Europe remains high even though the EMA restricted fluoroquinolone use in animals. However, flumequine, a quinolone similar to fluoroquinolones, is still used in livestock in the Netherland, Belgium, Greece or France. Therefore, we investigated whether flumequine selects for the same resistance mechanisms in *E. coli*.

To accomplish this, we investigated and enumerated resistant and non-resistant *E. coli* isolates obtained from caecal fermentation assays and from broilers exposed to different concentrations of flumequine and enrofloxacin.

Flumequine usage leads to approximately 3-fold increase of resistant *E. coli* in the caecal fermentation, comparable with enrofloxacin. After *in vitro* exposure to flumequine and enrofloxacin we detected the same SNPs (S83L, D87G) in *gyrA*. Furthermore, the same resistance-causing SNPs were found in phenotypic resistant *E. coli* isolates from broilers treated with enrofloxacin and flumequine.

Flumequine induces similar resistance mechanisms as enrofloxacin and restriction for use should be the same.

## Introduction

Antimicrobial compounds are arguably one of the most powerful discovered drugs in the history of human medicine ^1,2^. However, their success also seems to be their downfall since antimicrobial resistance (AMR) is a rising threat for the human and animal health ^3–5^. The usage of antimicrobials in livestock is contributing to the selection for AMR bacteria, as antimicrobials are often used metaphylactically, where the whole herd or flock is treated when only one or a few animals show illness ^6,7^. This large-scale use of antimicrobials in livestock can also select for AMR genes which can subsequently be transferred to other pathogens via horizontal gene transfer ^8^. The transmission of AMR and resistant bacteria via different pathways into the environment or to humans is a cause of concern ^6,9^. Therefore, it is crucial to have guidelines for antimicrobial resistance usage in husbandry to reduce the spread of AMR bacteria.

The European medicine Agency (EMA) has classified antimicrobial compounds in different categories from A (avoid) to D (prudence) to decrease usage, and parallel with that selection for AMR bacteria, of medically important antimicrobials for the human health ^10^. Although the EMA has classified all quinolones and fluoroquinolones among restrict usage the report of European Surveillance of Veterinary Antimicrobial consumption (ESVAC) in 2022 shows significant consumption of quinolones in Greece, The Netherlands, Belgium, and France ^11^. A reason for this could be caused by divergent categorization by the national commissions responsible for the antimicrobial usage in the veterinary practice. As a consequence, quinolones can be prescribed for livestock instead of individual animals since it is considered as a 2^nd^ choice antimicrobial compound in for husbandry usage in for example the Netherlands ^12^.

Quinolones are a widely used class of antimicrobials of which the first compound was nalidixic acid, synthesized by Lesher at al. 1962 ^13^. However, nalidixic acid was not used extensively because of the limited antimicrobial activity ^14^. Due to chemical modifications of the structure of quinolones, new drugs were created with greater potency, broader spectra of activity, improved pharmacokinetics and lower frequency of development of resistance ^14,15^. One of these chemical modifications led to the development of a class now known as fluoroquinolones. The resistance to (fluoro)quinolones can be related to three different mechanisms: (i) chromosomal single nucleotide polymorphism (SNPs) in the quinolone resistance determining region (QRDR), genes in this area include *gyrA, gyrB, parE* and *parC*. (ii) plasmid-mediated quinolone resistance (PMQR) genes encoded by *qnr* genes. (iii) upregulation of efflux pumps which can reduce drug accumulation ^16–18^.

All fluoroquinolones have the same mechanisms of action regardless of whether they are used in human or veterinary medicine; they inhibit the unwinding of the supercoiled DNA which enables the bacteria to perform protein synthesis and cell division. Even though none of the fluoroquinolones licensed for use in humans are approved for use in animals, there is still the possibility of increased fluoroquinolone resistance in human pathogens due to use of a veterinary approved quinolone. In the Netherlands there is an increasing trend in fluoroquinolone resistance in animals observed by the MARAN in species, such as *Salmonella, Campylobacter* and until 2022 also *E. coli* although it remains remarkably high ^19^. This is surprising since fluoroquinolones can only be administered to individual animals when there is no other alternative for individual usage. One explanation could be the use of flumequine which is classified as a second-choice antimicrobial, in contrast to enrofloxacin which is classified as a third-choice antimicrobial.

Considering the potential selection of flumequine for fluoroquinolone resistance, the main objective in this study was to investigate if flumequine has the same selective properties as enrofloxacin in *E.coli*. To establish this, we first conducted *in-vitro* work which consisted of a direct and a long-term exposure study of *E. coli* to concentrations of flumequine and enrofloxacin. Subsequently we studied the selection of fluoroquinolone resistant *E. coli* in a setting closer to reality by performing caecal fermentation assays treated with flumequine and enrofloxacin. Lastly, we sequenced *E. coli* isolates which we acquired from broilers treated with enrofloxacin or flumequine and investigated SNP occurrence in the QRDR region and phenotypic resistance. These experiments are covering *in vitro* and *in vivo* situations to investigate if the quinolone flumequine has the same selective properties as the fluoroquinolone enrofloxacin.

## Material and Method

### Obtaining fluoroquinolone resistant *E. coli* by direct exposure

The protocol on how to select resistant *E. coli’s* using ECOFF concentrations of fluoroquinolones was adapted from Cesaro et al., 2008 and Pasquali & Manfreda, 2007 ^20,21^. The parental strains 37 and 88 (supplementary data, File 1) obtained from fermentation assays from Swinkels et al. (2024), inoculated on blood agar plates and incubated overnight at 37 °C. One colony was taken from each plate and transferred to 10 ml LB broth and incubated overnight at 37 °C while shaking. Afterwards the cell cultures were centrifuged, and the pellet was suspended in 1 ml sterile LB broth to acquire an inoculum of ∼ 10^9^-10^10^ cfu/mL. Cell suspensions (100 µl) were inoculated on MacConkey agar plates containing the ECOFF concentrations of enrofloxacin (0.125 mg/L) or flumequine (2.0 mg/L), overnight at 37°C. Single colonies were taken from the selection plates and from both the enrofloxacin and flumequine resistant colonies. Afterwards the selected colonies were reinoculated on the opposite selection plate to determine co-resistance.

The MIC of the strains of interest were determined by standard broth microdilution adapted from EUCAST ESCMID (2003).

### Long-term exposure to fluoroquinolones with non-fluoroquinolone resistant *E. coli* strains

The long-term exposure experiment was derived and modified from Lenski et al. (1991) ^22^. For this experiment we selected 24 *E.coli* strains which all have been isolated from broilers and were susceptible for flumequine and enrofloxacin, except one positive control strain (supplementary data, file 2 (only the surviving strains)). Strains were stored in the −80 °C and therefore first inoculated on blood agar plates and incubated overnight at 37 °C. Subsequently, per strain a single colony was selected and transferred to a well containing 1 ml LB medium in a 12-wells plate (in total two 12-wells plate were used), referred to as T=0. The plates were incubated overnight at 37 °C, the next day 100 µl per strain was transferred to a well in a 96-wells plate. Additionally, with a stamp the strains were transferred to squared MacConkey plates containing flumequine 2 mg/L, enrofloxacin 0.125 mg/L and a control plate with no antimicrobials. This was done to determine the susceptibility of the 23 selected strain and the growth of the control strain. Next, 100 µl of the T=0 12-wells plates were transferred to a new 12-wells plate containing LB medium with a concentration of 0.0125 mg/L enrofloxacin (two 12-wells plates) and 0.2 mg/L flumequine (also two 12-wells plates), referred to as T=1. Subsequently the plates were incubated overnight at 37 °C and the next day the strains were transferred to squared McConkey selections plates with ECOFF concentrations of flumequine and enrofloxacin. This procedure was repeated for 10 days with each day an increase in the concentrations, 0.0125 mg/L for enrofloxacin and 0.2 mg/L for flumequine. Every day the proportion of non-wildtype (referred to as resistance) at ECOFF concentrations was determined. After 10 days the strains were exposed to clinical breakpoint concentrations of enrofloxacin (2 mg/L) and flumequine (8 mg/L) and analysed for survival and resistance.

### Caecal fermentation to determine co-resistance of *E. coli* from cecum faeces

The protocol for the fermentation studies was completely adapted from Swinkels et al. (2023) ^23^. Only alteration or addition was the use of flumequine (2 mg/L, 0,2 mg/L and 0.02 mg/L). The caecum material was obtained from broilers from commercial farms in the Netherlands which encountered serious lameness. The caeca were removed from euthanized broilers which were used for educational purposes at the faculty of veterinary medicine (registration-number: AVD10800202115056). The broilers aged between 15 to 40 days, due to different sample moments as this experiment was repeated three times and for every experiment caecum material of two different broilers was used.

### Resistance characterization of *E. coli* isolates from poultry farms treated with clinical concentrations of flumequine

Samples, in terms of cloaca swabs, were taken from commercial poultry farms when a treatment with flumequine needed to be performed according to a veterinarian. In total 16 broilers were swabbed in the poultry house before treatment and 16 in the untreated poultry house as a baseline measurement. After the treatment again 16 broilers were swabbed in the untreated and treated poultry house. The cloaca swabs were transferred to the lab for further processing at the sampling day. In the lab the swabs were first inoculated on MacConkey plates and incubated overnight at 37 °C. Afterwards two colonies (per swab) from each MacConkey plate were transferred to blood agar plates and incubated at 37 °C overnight. Subsequently, the colonies, 32 in total per timepoint and treatment, were transferred to separate wells in a 96-wells plate from which enabled transferring the colonies to squared MacConkey plates. The squared MacConkey plates were containing concentrations of flumequine 2mg/L (ECOFF), 8 mg/L (clinical breakpoint) or enrofloxacin 0.125 mg/L (ECOFF), 2 mg/L (clinical breakpoint) and a control plate without any antimicrobials. After overnight incubation the proportion of resistant *E. coli* could be calculated with comparing the selection plates to the control plate.

### Sequencing of *E. coli* isolates

Several experiments were followed up by sequencing for SNP and sequence type determination. For that reason, DNA extraction of *E. coli* was preformed using the NGS DNeasy UltraClean Microbials kits from Qiagen. Additionally, the concentrations of all the DNA isolates were checked using Thermo Fishers’ Qubit Fluorometer to guaranty high enough concentration for sequencing.

#### MinION Sequencing

Sequencing was done following the Oxford Nanopore MinION using the Rapid barcoding Kit (SQK-RBK096) and sequenced in R9.4.1 flow cells (FLO-MIN106) using the MinION Mk1B (ONT) device. ONT raw reads were subjected to base-calling using MinKNOW (v4.5.3) with the Super Accurate model. Afterwards, the reads were trimmed with Filtlong v0.2.1 and the quality was assessed with FastQC (v0.11.4). Reads were assembled to contigs with Flye v2.9. Assemblies were polished using Medaka 1.4.3 (https://github.com/nanoporetech/medaka) and Homopolish 0.3.4. The quality of all sequences was checked with Checkm v1.1.3, and only genomes with a contamination threshold of <5% and completeness threshold of >95% were included in the analysis. ResFinder v4.0 was used to identify the SNPs and MLST was determined using mlst.

#### Illumina sequencing

Illumina sequencing was performed using Illumina Nextseq 500 (Useq, Utrecht sequencing facility) with a maximum read length of 2 x 150 bp. Libraries were prepared with Illumina Nextera XT DNA Library Preparation Kit according to the manufacturers protocol ^24^. Read processing and assembly was performed as described for Illumina reads in Swinkels et al. 2023 ^23^.

All sequences can be found in the SRA, Supplementary data.

## Results

### *gyrA* mutations in *E. coli* after direct exposure to ECOFF concentrations

At first, we conducted a direct exposure on selection plates with fluoroquinolone and enrofloxacin to study if we could obtain resistant isolates and determine co-resistance (Table 1). The isolates were collected as single colonies from selection plates, and we further characterized the strains by phenotypic resistance, MIC levels and SNP presence in *gyrA* gene. After sequencing, we found the SNP S83L is mostly occurring after inoculation with enrofloxacin while flumequine selects more often for the D87G SNP but also for the S83L SNP. Furthermore, we found that a SNP in S83L or D87G results in growth on selection plates with enrofloxacin 0.125 mg/L and flumequine 2 mg/L. Contrasting is the observation that the MIC for both enrofloxacin and flumequine is higher in the isolates with the S83L SNP compared to the isolates harbouring the D87G SNP.

**Table 1.**
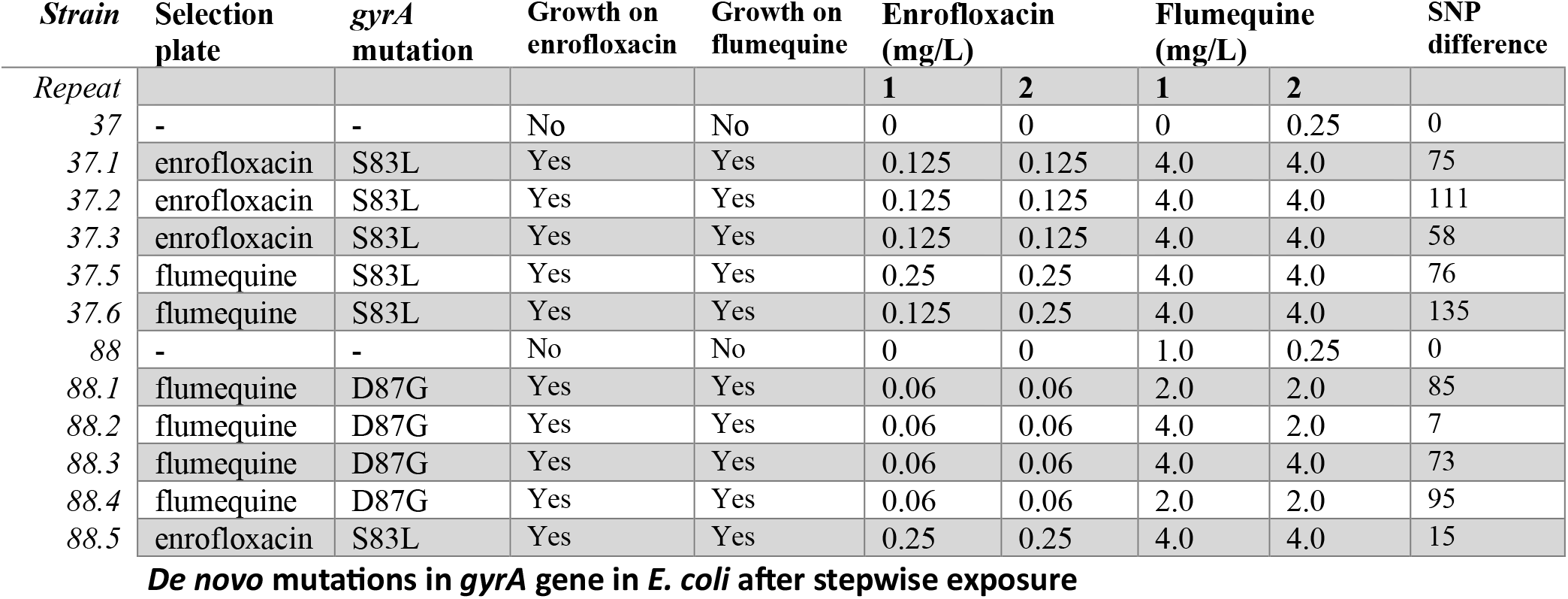
Characterization of isolates from direct exposure. In the columns the SNPs in the GyrA gene, phenotypic resistance, MICs of flumequine and enrofloxacin are displayed. The last column shows the SNP difference between the parental strain and the obtained isolates.

### *De novo* mutations in *gyrA* gene in *E. coli* after stepwise exposure

After the direct exposure on selection plates, we performed a stepwise increasing exposure to susceptible *E. coli* isolates to observe if induction of *de novo* mutations over time in the *gyrA* was possible. We started with 24 strains, however after MinION nanopore sequencing we had to exclude 14 strains due to different sequence type at T=9. In Figure 1 we can observe the increased resistant strains after treating for enrofloxacin as well for flumequine, 100% proportion resistance is reached for either the flumequine or enrofloxacin treatment. Interestingly, 100% flumequine resistance is reached before 100% enrofloxacin resistance is occurring in the enrofloxacin treatment. This phenomenon is also observed when treating with flumequine when 100% resistance towards enrofloxacin is reached before 100% flumequine resistance. Nevertheless, we can clearly observe co-resistance between enrofloxacin and flumequine while treated with one of these antimicrobials. Furthermore, the low concentrations of either enrofloxacin or flumequine can already induce a phenotypic resistance at ECOFF concentrations.

**Figure 1.**
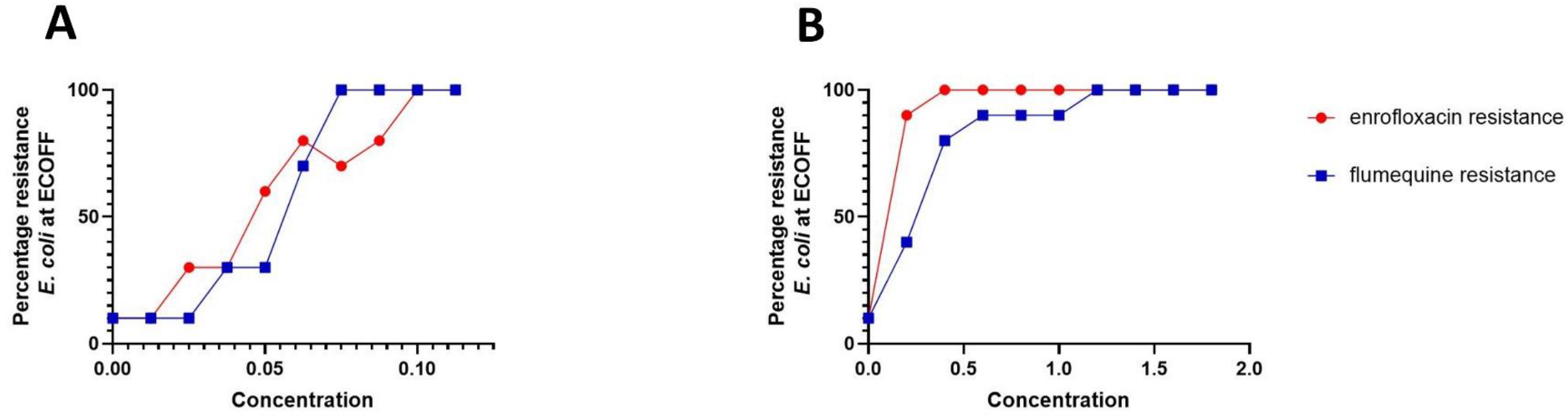
Percentage of surviving E. coli isolates after stepwise exposure. **A** represents the curve after the enrofloxacin treatment and **B** after the flumequine treatment.

After MinIOn nanopore sequencing, we determined the sequence type of the *E. coli* strains at T=9 in combination with chromosomal SNPs in het QRDR region. We observed (Table 2) that the sequence type of the strains at T=9 was identical at T=0 of the experiment and in combination with one or more SNPs occurring at T=9 shows the occurrence of *de novo* mutations. For some strains a sequence type was not determined but with a spanning tree we could observe clustering of the different strains which allowed us to see their phylogenetic relatedness (supplementary data). In Table 2 we can also observe that some strains did not show any point mutation which should enhance their resistance towards enrofloxacin or flumequine. Nevertheless, these strains showed resistance as we can observe in Figure 1.

**Table 2.** Point mutation in QRDR and sequence type (ST) of the E. coli strains at the start of the long-term exposure T=0 and after treatment with flumequine or enrofloxacin at T=9. * SLV=Single Locus Variant, ** DLV= Double Locus Variant.

|  | **T=0** |  | **T=9**<br>**flumequine** |  | **T=9**<br>**enrofloxacin** |  |
| --- | --- | --- | --- | --- | --- | --- |
| *Strain* | *Point*<br>*mutation* | *ST* | *Point*<br>*mutation* | *ST\** | *Point*<br>*mutation* | *ST\*\** |
| WGS-ecoli-sample-38 | - | 5334 | *gyrA*<br>p.A119E | 5334 | Unknown | 5334 |
| WGS-ecoli-sample-46 | - | 349 | *gyrA* p.D87H | 349 | *gyrA* p.D87G | 349 |
| WGS-ecoli-sample-47 | - | 8070 | *gyrA* p.D87Y | 8070 | *gyrA*<br>p.D678E | 8070 |
| WGS-ecoli-sample-49 | - | 58 | *gyrA* p.D87G | 58(SLV)* | *gyrA* p.D87G | 58 |
| WGS-ecoli-sample-64 | - | 4642 | *gyrA* p.D87G | 4642 | *gyrA* p.D87G | 4642 |
| WGS-ecoli-sample-70 | - | 2307 | *gyrA* p.D87G | 2307 | *gyrA* p.D87G | 2307(DLV)** |
| WGS-ecoli-sample-88 | - | 40 | *gyrA*<br>p.L447M | 40 | *gyrA* p.D87G | 40 |
| WGS-ecoli-sample-107 | - | 101 | *gyrA*<br>p.E153G | 101 | *gyrA* p.D87G | 101 |
| WGS-ecoli-sample-113 | - | 720 | *parC* p.S57T | 720(SLV)* | *gyrA* p.D87Y,<br>*parC* p.S57T | 720 |
| WGS-ecoli-sample-99 | *gyrA* p.S83L | 752 | *gyrA* p.S83L |  | *gyrA* p.S83L | 752 |

### Quantifying fluoroquinolone resistant E. coli proportions from caecal fermentations

After conducting *in vitro* experiments which showed that both de novo mutations and selection for existing fluoroquinolone resistant *E. coli* was possible, we aimed our focus to a more natural setting by performing caecal fermentation with fresh caeca material. The caecal fermentation assays provided a clear result in terms of selective advantages for resistant strains in a more complex setting which harbours natural *E. coli* strains, Figure 2. The highest concentrations of flumequine 2 mg/L and enrofloxacin 0.125 mg/L showed an increase in the resistant *E. coli* strains. More interestingly, the proportions in both caecal fermentation assays, treated with flumequine or enrofloxacin, showed similar counts for resistance against flumequine and enrofloxacin. The treatment with enrofloxacin 0.125 mg/L showed an almost 3-fold increase in the number of resistant isolates for flumequine or enrofloxacin compared to the control group at 6 or 30 hours. The same result we observed in the caecal fermentation treated with flumequine 2 mg/L, a 3-fold increase in number of resistant isolates in flumequine and enrofloxacin compared to the control group 6 or 30 hours.

**Figure 2.**
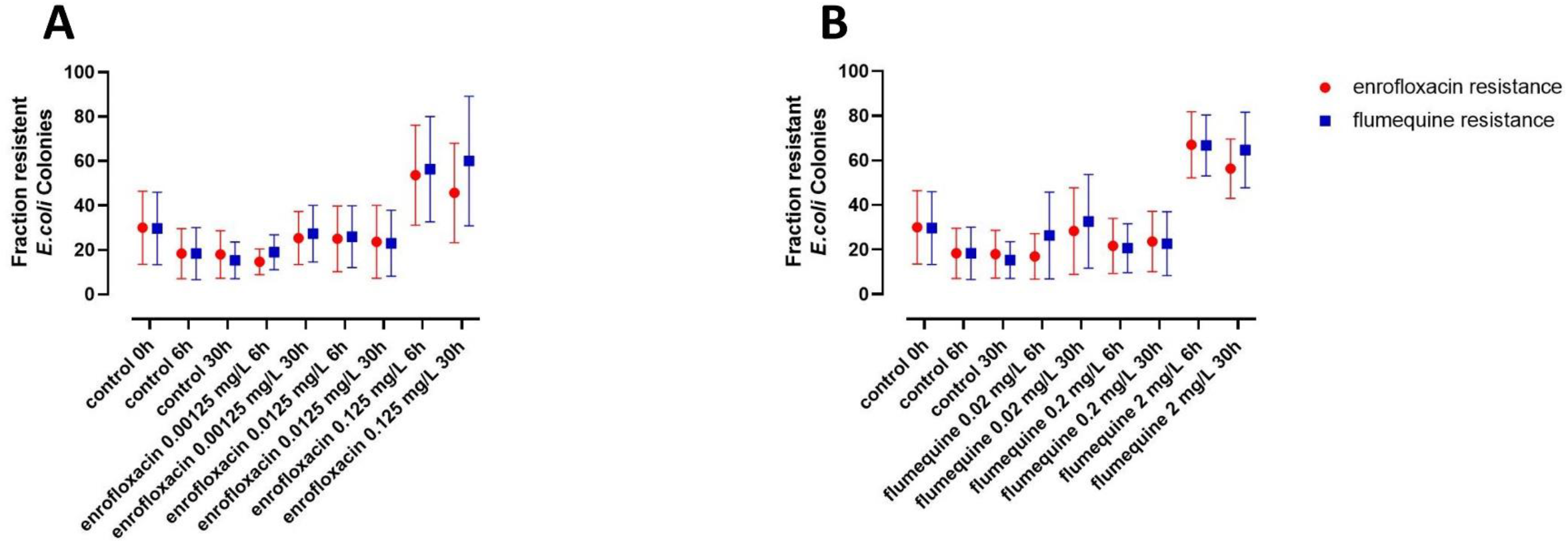
Resistant E. coli colonies isolated from the caecal fermentations. Graph **A** the caecal fermentation treated with enrofloxacin and graph **B** the caecal fermentation with flumequine. Error bars are representing standard errors of the mean.

### Fluoroquinolone resistant *E. coli* after therapeutic exposure to flumequine

Cloaca swabs from broilers after therapeutically administrated with flumequine were screened for enrofloxacin and flumequine resistant *E. coli*, as is shown in Figure 3. The control group showed no remarkable differences of *E. coli* isolates growing on selection plates with flumequine or enrofloxacin between T=0 and T=1. On the other hand, in the flumequine treatment group we observed almost a 4-fold increase of ECOFF (flumequine and enrofloxacin) resistant *E. coli* isolates at T=1 compared to T=0. We even observed a small increase in clinical breakpoint resistant *E. coli* isolates at T=1 after treatment, for either flumequine and enrofloxacin. Although this increase seems not as substantial as the increase at ECOFF concentrations, it is still around 3.5-fold more compared to T=0. In comparison with therapeutic concentrations of enrofloxacin we observed similar co-resistance with flumequine, supplementary File 3.

**Figure 3.**
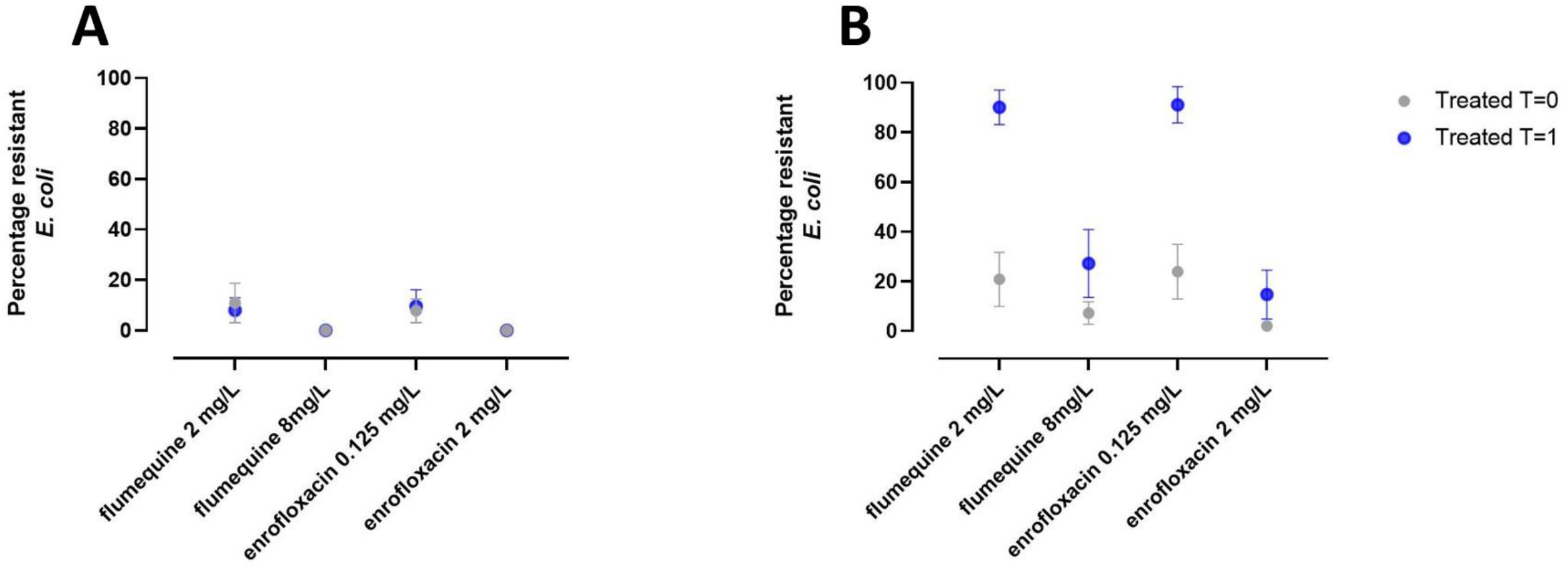
Percentage resistant E. coli colonies after exposure of clinical concentrations of flumequine. T=0 is before initiating the treatment and T=1 directly after termination of the treatment. **A** represents the control group which was not treated with flumequine and **B** the group that was treated with flumequine. On the x-axis the concentrations in the control plates are displayed. ECOFF concentration (2 mg/L flumequine and 0.125 mg/L enrofloxacin) and clinical breakpoint concentrations (8 mg/L flumequine and 2 mg/L enrofloxacin).

Furthermore, we sequenced with 32 susceptible isolates from the treated group at T=0 and 16 *E. coli* isolates which have shown phenotypic growth on clinical breakpoint concentrations of enrofloxacin after treatment, shown in Table 3. We observed that the isolates from the clinical breakpoint selection plates had three SNPs in the QRDR (*gyrA* p.S83L, p.D87N, *parC* p.S80I) and mostly had the same sequence type. We compared the sequence type of resistant *E. coli* after treatment with the sequence types of the isolates sequenced at T=0 but we were not able to find a susceptible *E. coli* strain with the same sequence type. We could therefore not specifically observe inducing of the SNPs as a consequence of flumequine treatment. However, we did observe selection for enrofloxacin clinical breakpoint resistant *E. coli* after treatment with clinical concentrations of flumequine.

**Table 3.**
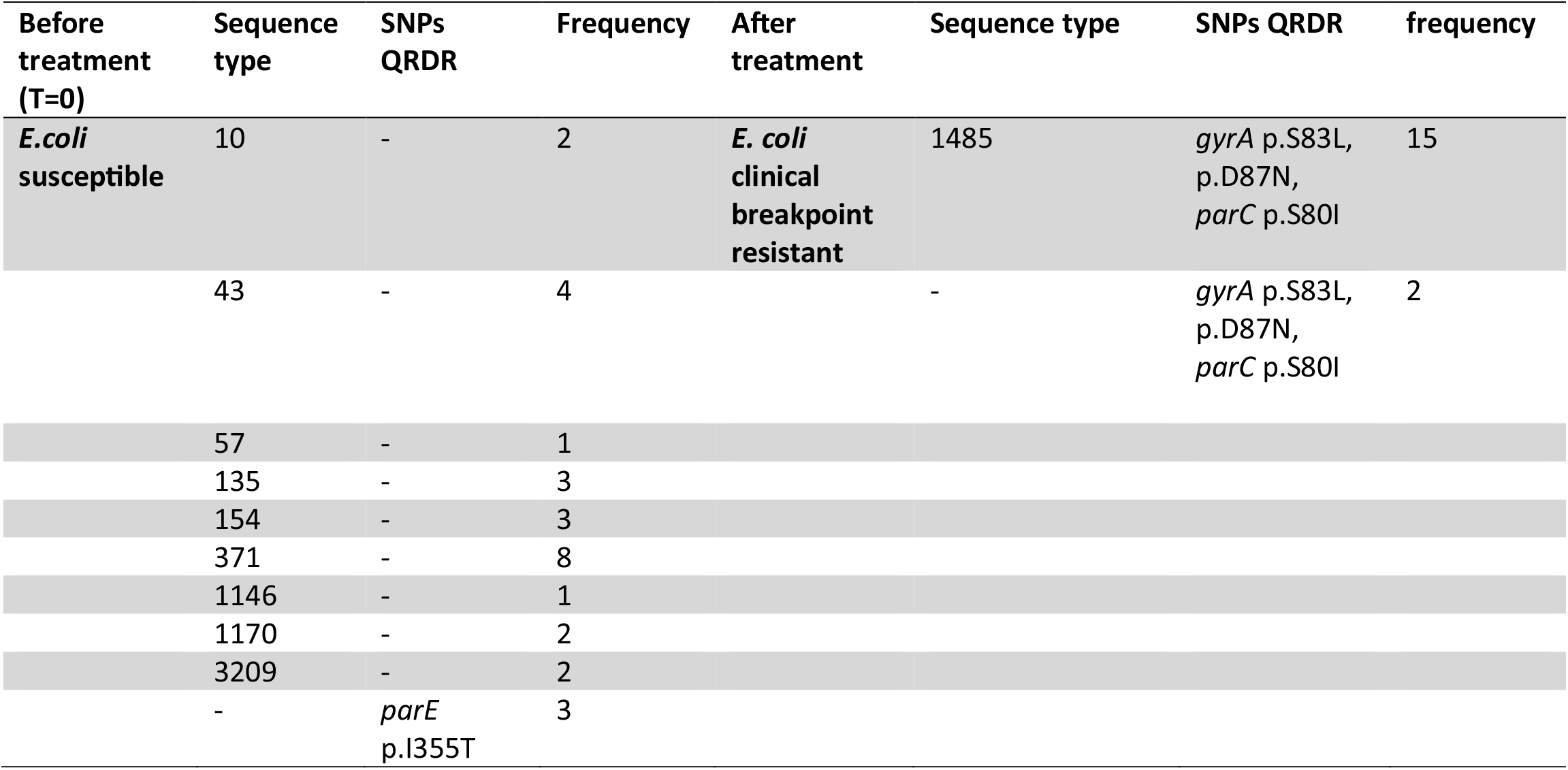
Characteristics of E. coli isolates before and after flumequine treatment with therapeutical concentrations. In the columns the sequence type, SNPs and the frequency of the isolates are displayed.

| Before treatment (T=0) | Sequence type | SNPs QRDR | Frequency | After treatment | Sequence type | SNPs QRDR | frequency |
| --- | --- | --- | --- | --- | --- | --- | --- |
| *E.coli* susceptible | 10 | - | 2 | *E. coli* clinical breakpoint resistant | 1485 | *gyrA* p.S83L, p.D87N, *parC* p.S80I | 15 |
|  | 43 | - | 4 |  | - | *gyrA* p.S83L, p.D87N, *parC* p.S80I | 2 |
|  | 57 | - | 1 |  |  |  |  |
|  | 135 | - | 3 |  |  |  |  |
|  | 154 | - | 3 |  |  |  |  |
|  | 371 | - | 8 |  |  |  |  |
|  | 1146 | - | 1 |  |  |  |  |
|  | 1170 | - | 2 |  |  |  |  |
|  | 3209 | - | 2 |  |  |  |  |
|  | - | *parE* p.I355T | 3 |  |  |  |  |

## Discussion

Flumequine is selecting for fluoroquinolone resistant bacteria. Flumequine specifically selects for bacteria which acquired a mutation in the QRDR region, in genes coding for *GyrA* or *ParC* genes to be precise. This mutation can be pre-existing or can occur de novo during treatment. The usage of flumequine for treating livestock in several European countries occurs for example in the Netherlands and Greece (≥0.75 mg per population correction unit) according the ESVAC ^11^. This is concerning as its potentially increases fluoroquinolone resistant bacteria within livestock which is not only a risk for treatment failure of animals but also for the possible transfer of these AMR resistant bacteria to humans. Fluoroquinolones are an important class of antimicrobials for treating infections in human medicine ^25^. Fluoroquinolones such as ciprofloxacin, levofloxacin or norfloxacin will no longer be effective due to a similar or identical resistance mechanism as is induced or selected by using flumequine ^26^.

In this research we mainly focused on the ECOFF concentrations to distinguish the *E. coli* isolates from wildtype and non-wildtype. This concentration is used mainly for surveillance purposes ^27^. If growth was observed, the *E. coli* isolate was scored as resistant. One can argue that this is not clinically relevant. However, if the *E. coli* isolate can grow on ECOFF concentrations it is reasonable to expect that mutations are acquired resulting in resistance.

We encountered enrofloxacin clinical breakpoint resistant *E. coli* isolates after treatment of broilers with clinical dosages of flumequine. This is underlining that flumequine is able to select for fluoroquinolone clinical breakpoint resistant bacteria when used for farm animals. We did not demonstrate that flumequine used in the farm environment can induce SNPs in the QRDR leading to fluoroquinolone resistance, likely because of the high diversity of strains in a broiler flock and the difficulty of obtaining the same clone before and after treatment. However, we did prove that a single SNP in *E. coli* can be induced by the usage of flumequine and these isolates are able to withstand ECOFF concentrations of either flumequine or enrofloxacin. In terms of clinical breakpoint resistance mostly two or multiple SNPs in the QRDR are necessary ^18,28^. This means that the mutation frequency towards clinical resistance in the QRDR region is much lower if there is already one SNP present in the QRDR. If the mutation frequency is 10^−6^ per cell division it means that two SNPs in QRDR region may occur in 10^−12^ cell divisions and when one SNP is already present the potential to get clinically resistant *E. coli* is 10^−6^ cell divisions less, hugely increasing to probability for clinical breakpoint resistant *E. coli* ^29,30^.

We used the indicator organism *E. coli* in this research to determine whether flumequine had similar inducing and selecting properties as enrofloxacin. *E. coli* is widely used as an indicator organism for resistance levels as a consequence of the ease it can be propagated in the lab and studied due to its well-known genomics ^31^. Still *E. coli* is just a single bacterial species in the very extensive microbial species composition, for example 15% in the faeces of healthy broilers ^32^. Therefore, it might be interesting to study the SNP occurrence of flumequine in other species and also observe if this is also enhancing and spreading fluoroquinolone resistance. For example, in the Netherlands fluoroquinolone resistance in *Campylobacter* is increasing ^19^, which might be due to flumequine usage.

The ESVAC has classified quinolones in two subgroups; fluoroquinolones and other quinolones. Among other quinolones we encounter flumequine, oxolinic acid and cinoxacin. This divergent classification might be arbitrary if we consider the outcome of our research, in which we clearly show that flumequine has selective properties which enhance fluoroquinolone resistance. As a result, it would be interesting to study the effect of oxolinic acid or cinoxacin on the selection for SNPs in the QRDR of chromosomal *gyrA* and *parC* genes. Especially since oxolinic acid is used in aquaculture where it can be potentially a factor to increase fluoroquinolone resistance ^33–35^. A study conducted by Ham et al. (2022) showed that oxolinic acid used in apple and pear orchards can also select for SNPs in the QRDR of *Erwinia amylovora* ^36^. This is implying that oxolinic acid, described as another quinolone, has the same effect as flumequine and to elaborate oxolinic acid is a non-fluorinated quinolone in contrast to enrofloxacin and flumequine.

In the stepwise exposure experiment we observed some peculiar results which were unexpected. First, we observed SNPs in *gyrA* in the *E. coli* isolates at T=9, after treatment with either flumequine or enrofloxacin. However, some SNPs were not included in the Resfinder database and were recently discovered as SNPs in the QRDR inducing resistance ^37^. Among these SNPs we found the following; *gyrA* p.A119E, *gyrA* p.D678E, *gyrA* p.L447M and *gyrA* p.E153G ^38–40^. Furthermore, we encountered one strain, WGS-sample-38, which showed phenotypic resistance after treatment with enrofloxacin, but we did not detect a corresponding SNP. This could attribute to a SNP which is not known to induce fluoroquinolone resistance or that efflux pumps were upregulated during the experiment ^41^. Secondly, there was a rapid increase in the percentage of resistant *E. coli* at ECOFF concentrations even when exposed to very low concentration of enrofloxacin or flumequine. This underlines that low concentrations of (fluoro)quinolone can already induce resistance which illustrate the effect of residual antimicrobial concentrations ^9,42^. Finally, we observed in susceptible *E. coli* isolates at T=0 from the field samples, that some *E. coli* harboured a SNP in the *parE* gene that also belongs to the QRDR. Although this feels counterintuitive since the *E. coli* isolates were scored susceptible, but in the literature, it is shown that a single mutation in the *parE* gene does not result in growth on ECOFF concentrations ^43^, however this could lead to higher resistance when combined with other mutations.

Given the selection of flumequine among the *E. coli* isolates in this study, authorities regulating or setting the prescription of antimicrobials for livestock should abolish the differentiation between fluoroquinolone and “other quinolones”. Especially since flumequine can create a reservoir of fluoroquinolone resistant bacteria which can transfer to humans or the environment ^44^. Institutions as EMA already classified all quinolones among restricted usage which is according to our results appropriate to reduce AMR resistance ^10,11,45^. However, national advisory groups such as the Dutch WVAB allow flumequine as a 2^nd^ choice antimicrobial (can be used after 1^st^ choice) instead of a 3^rd^ choice antimicrobial (restricted and only for individual animals) ^46^. This classification might explain the relatively high fluoroquinolone resistance in The Netherlands in broilers ^19^. Moreover, the European Centre for Disease Prevention and Control (ECDC) reported in 2023 that fluoroquinolone resistance in *E. coli* isolates from humans is above 25 % in 17 of the 45 countries ^47^. This emphasizes that fluoroquinolones resistance remains a multi sectoral problem.

To conclude, flumequine which shows the same selective properties as enrofloxacin, should be considered as fluoroquinolones for the usage in livestock as this would have potential impact on the reduction of AMR bacteria.

## Acknowledgements

This work was supported by the Netherlands Centre for One-Health (NCOH). We also would like to thank Jeroen Leus for helping collect the field samples. Furthermore, we would like to thank the suppliers of the caecal material which have been used in this study.

## Ethics declaration

Competing interests

The authors declare no competing interests.

## Supplementary information

Sequence data for this article can be found in the SRA under accession PRJEB75157.

Supplemental File 1, XLSX file, 10 kB

Supplemental File 2 (only surviving strains), XLSX file, 10 kB

Supplemental File 3, PDF file, 334 kB

